# Global connectivity and networks of marine reserves

**DOI:** 10.1101/2022.11.20.515214

**Authors:** Julia McDowell, Marco Andrello, Laure Velez, Nicolas Barrier, Stephanie Manel, Laura J. Pollock, David Mouillot

## Abstract

Cooperation between countries in managing and protecting shared marine resources is beneficial both ecologically and economically, but how best to establish the cooperation needed at a global scale is largely unknown. Here, we used hydrodynamic modelling to identify ecologically connected networks of marine reserves (MRs) and evaluated these networks with socio-economic indicators. Most of the networks are homogenous with similar levels of development, shared languages, and other cultural values. However, we found that 17% (11/66) of the largest networks (>20 MRs) span multiple countries. These heterogenous networks are composed of countries with different economic, political, and cultural views. Countries that control more, larger marine reserves also have a more even ratio of source reserves to sinks. We discuss that, while such economic and cultural homogeneity might lead to more efficient ecological management in the short term, heterogeneous networks may prove to be more resilient in the longer term, when climate change will modify marine connectivity.

## 1 Introduction

Marine ecosystems are highly threatened by anthropogenic pressures such as overfishing, repeated heat waves, and overuse (Hughes et al. 2019, O’Hara 2021). Marine protected areas (MPAs) are an effective way of mitigating some of these pressures (Roberts 2017, Duarte et al. 2020). MPAs restrict human use to protect both living and non-living resources making them useful management tools (Edgar 2014). Especially effective are marine reserves (MR), which are no-take MPAs (Costello & Ballantine 2015) that allow for larger fish and greater population sizes both within their boundaries and in the close vicinity (Edgar 2014, Ohayon 2021). In addition to the well-documented ecological and fisheries benefits, marine reserves can also promote tourism, human wellbeing, and alternative livelihood strategies (Ban et al. 2019). Given the potentially large benefits of marine reserves, it is important to understand which factors contribute the most to reserve success. There is evidence that successful reserves (e.g. those that have greater fish biomass relative to unprotected areas) have high compliance to fishing restrictions, are older, larger, and isolated from humans (Edgar et al. 2014; Cinner et al. 2018). What is less known is the impact of the greater network of reserves on effectiveness.

There are many reasons that the connectedness of reserves could be beneficial from a conservation perspective. Connectivity promotes source-sink dynamics, gene flow, demographic and genetic rescue, and also assists in species range shifts (Andrello et al. 2017; Roberts, 2017; Manel et al. 2019). Isolated marine reserves are slow to recover after disturbance events; the isolation impeding larval recruitment and significantly slowing recovery time (Olson 2019). Conversely, oceanic currents also mediate the spread of marine pollution, invasive species, and diseases (Jönsson & Watson 2016; Jaspers et al 2018). Therefore, hydrological connectivity among MRs across the seascape is a key process capable of modifying the functioning and the effectiveness of MR networks. Further, understanding how reserves are hydrodynamically connected is necessary to promote long-term ecological and socio-economic resilience under ever increasing disturbances; reserves designed with optimized connectivity in mind greatly improve the benefits of the reserve (Edgar 2014, Sala 2018, Goetze 2020). For these reasons, the Convention for Biological Diversity, through the post-2020 global framework for biodiversity current under discussion, recognizes that connectivity is a necessary component of MR networks and needs to be included in their design (https://www.cbd.int/conferences/post2020), and oceanographic connectivity is increasingly considered in spatial conservation planning (Maina 2019).

So far, this progress in understanding how connectivity impacts marine reserves has mostly been ecological in nature. Understanding how political, economic, and societal collaboration among countries align with the ecological connectivity among their marine reserves at a global scale is largely unknown (Treml et al 2015). It would be unfair to assume every country is capable of the same level of involvement in implementing new protected areas and reserves due to the high cost, and so managing connected networks will necessitate an understanding of the “burden” that each country is capable of sustaining (Maina 2019). Previous studies have shown that hydrodynamically connected networks of MRs can span Exclusive Economic Zones (EEZ) of multiple countries that have different standards of living with varying levels of governance, environmental performance, and dependency on marine resources (Andrello et al. 2017; Wendling et al 2020). Managing and conserving connected ecosystems requires increasing levels of political and cultural connectedness and collaborations across scales and jurisdictions (Bodin 2016).

A broad coalition of environmental organizations are campaigning to protect at least 30% of the amount of land and sea by 2030 (Dinerstein 2019), but this effort could be hindered by our limited understanding of how countries interact within ecologically connected networks of MRs on a global scale. Countries that have a desired outcome in common, be it increased wealth, better environmental protections, or another goal should have an easier time creating policies that can be agreed upon by all parties involved. By contrast, countries with different ideas or with different dependencies upon marine areas for subsistence are less likely to reach common agreements regarding the levels of protection and management of marine areas (Treml 2015).

Here, we aim to document the large-scale connectivity between marine reserves and their degree of isolation while also characterizing the social configuration of reserve networks. We first identify all the hydrodynamically connected networks of marine reserves in the world, as well as the highly isolated reserves. We then identify the most well-connected countries, as well as countries that have significantly higher ratios of incoming or outgoing connections. We last describe the shared cultural, political, and economic characteristics of the countries which control the largest networks, in an attempt to understand why they are so successful in creating highly connected, multi-country networks.

## 2 Materials and Methods

### 2.1 Marine protected areas and marine reserves

The MPA database was downloaded from the World Database on Protected Areas and filtered to only keep protected areas known as ‘marine’ (WDPA, downloaded January 2019). From there we retained only the MRs that are either in IUCN category Ia or Ib or designated as at least partially no-take, since only marine reserves provide widely recognized ecological benefits (Turnbull 2021). The final database includes 2,203 marine protected areas that we refer to as marine reserves (MRs) in the following sections.

### 2.2 Hydrodynamic connectivity between marine reserves

We evaluated hydrodynamic connectivity using Lagrangian models of dispersal. We used sea surface current velocities from Copernicus Marine Environment Monitoring Service from January 2008 to March 2017. The horizontal resolution of the current velocity datasets was 1/12^th^ degree, and the temporal resolution was one day. We released 10,000 virtual particles from the centroid of each marine reserve and tracked their trajectories using Ichthyop 3.3 (Lett et al. 2008). This was designed to show the trans-boundary dispersal, and not capture local retention and higher variability that could been seen with a higher resolution.

We calculated the probability of connection between all pairs of MRs. The spatial position of particles relative to MR polygons was assessed using the function ‘gContains’ in the R package rgeos 0.3–19 (Bivand & Rundel, 2017) on the latitude and longitude of each particle. Connection probabilities were used to construct a connectivity matrix among all MRs. Each value is the probability that particles released in one marine reserve are recruited in another, thereby giving the probability of inputs and outputs of each marine reserve to and from all others. Networks are a set of connected MRs, and were determined using the ‘clusters’ function of the R package igraph 1.0.1 (Csardi 2006). However, direction of the connection was disregarded to create the networks.

Networks were initially found for multiple pelagic particle durations, including 20, 40, and 60 days (Appendix 2, 3). Ultimately, we chose to analyze based on a 40-day particle duration because it is closest to the mean pelagic duration of fish larvae (Booth and Parkinson 2011, Graham et al. 2008, Jones et al. 2009).

We performed analyses at the network and at the country level. At the network level, we analyzed the indicators for countries involved in each individual network. At the country level, we analyzed the connections that the MRs controlled by each country have to other countries. More detailed methods are available in Appendix 1.

### 2.3 Social, economic, and political indicators

We used the fisheries dependency index and the human development index to characterize the socioeconomic dimensions of MR networks. The fisheries dependency index assesses the importance of a coastal country’s fisheries in terms of their contributions to a national economy, employment, and food security (Andrello et al. 2017). The fisheries dependency index is bounded between 0 and 100, with higher values indicating higher dependency. We used the 2018 Human Development Index (HDI) (United Nations Development Program, downloaded 2019) as an indicator of human development. The HDI is based on national statistics of life expectancy, education level, and gross national income per capita, ranges from 0 to 1, with higher values indicating higher development.

We used the 2018 Environmental Performance Index (EPI) as an indicator of the importance each country places on the environment (Wendling et al 2020). The EPI ranks 180 countries on two dozen indicators spanning ten categories across environmental health and ecosystem vitality, including air pollution, climate and energy, and fisheries. The EPI is produced jointly by Yale University and Columbia University in collaboration with the world economic forum, ranges from 0 to 100, with higher values indicating higher importance placed on environmental issues.

Data on languages were taken from the Ethnologue Global Dataset compiled by SIL International, a comprehensive database of all languages present in each country (Eberhard 2019). It also documents the number of speakers, literacy rate, and characteristics of each language. In this study, we used the official language of the country, either *de facto* or *de jure*, in which language governmental and cross border issues would be discussed. If a country has two official languages, both were included in the analysis.

## 3 Results

### 3.1 The global structure of marine reserve networks

We identified a total of 65 networks of marine reserves based on a cluster analysis. Out of 2,203 marine reserves described at the time of the study, 1,618 (73.4%) are included in a network of more than one reserve. Of the 65 total networks, we further focused on the largest 11 networks that each are composed of 20 or more marine reserves (Fig. 1). These 11 networks contain 60.7% (1339) of marine reserves as well as 42.1% of the total surface area of all marine reserves.

**Figure 1:**
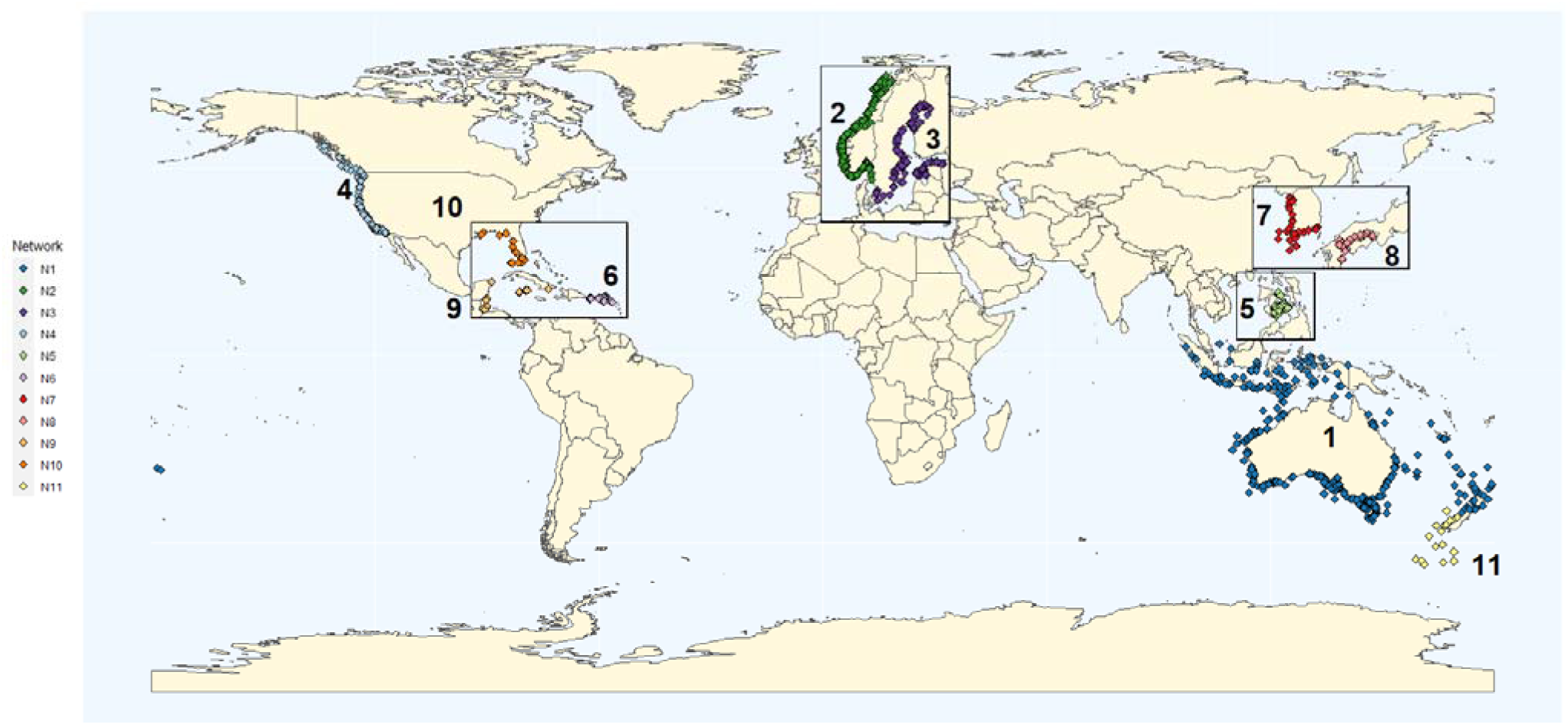
The 11 largest networks of marine reserves (with 20 or more connected reserves) based on a 40-day pelagic larval duration. See appendix 4 for detailed information on the country and language composition of each network.

The largest network in terms of the number and size of reserves links the Indian and Pacific oceans (N1, Fig. 1, Appendix 4). N1 has 475 reserves, spans five countries, and covers more than twice the area of the next largest reserve network. The second largest network in terms of the number of reserves is in the Norwegian Sea with 374 reserves, composed primarily of Norwegian reserves with a few Swedish reserves (N2, Fig. 1). Despite its high quantity of reserves, N2 is much smaller than N1 in terms of size, ranking 43rd by area covered out of all 66 networks. The other nine top networks range in size from 20 to 200 reserves. The third largest network is located in the Baltic region and three large networks are located in the Caribbean—one in the eastern Caribbean (N6), one in the western Caribbean (N9) and one in the Gulf of Mexico (N10). By contrast, Africa, South America, and Eastern Indian Ocean have no networks that rank in the top 11, despite each having multiple marine reserves. Similar to the number and size of reserves in each network, the number of involved countries is also variable ranging from multi-national networks spanning five countries (N1 in and around Australia and N6 in the easter Caribbean) to networks completely within the domain of a single country. Of the total 66 networks identified, 51 of them are single nation networks.

While most marine reserves are connected to at least one other reserve, 585 (26.6%) out of the analyzed reserves are not connected to any single other reserve. These isolated reserves are widespread (Figure 2), even in areas that are regionally dominated by large networks. For example, Australia, Norway, and Sweden are all surrounded by isolated marine reserves while at the same time being the locations of the top three largest networks. These isolated reserves are not always small; isolated reserves are some of the largest by area with eight of them each covering 10,000 square kilometers. For example, the South Georgia and South Sandwich Island marine reserve is over 1.24 million square kilometers, yet has no probability of connectivity to other sources. However, most isolated reserves (394, or 57.9%) are less than 1 square kilometer and 490 (78.0%) are under 10 square kilometers. Many of the smallest marine reserves are located in Norway and Sweden, an area of generally high connectivity.

**Figure 2:**
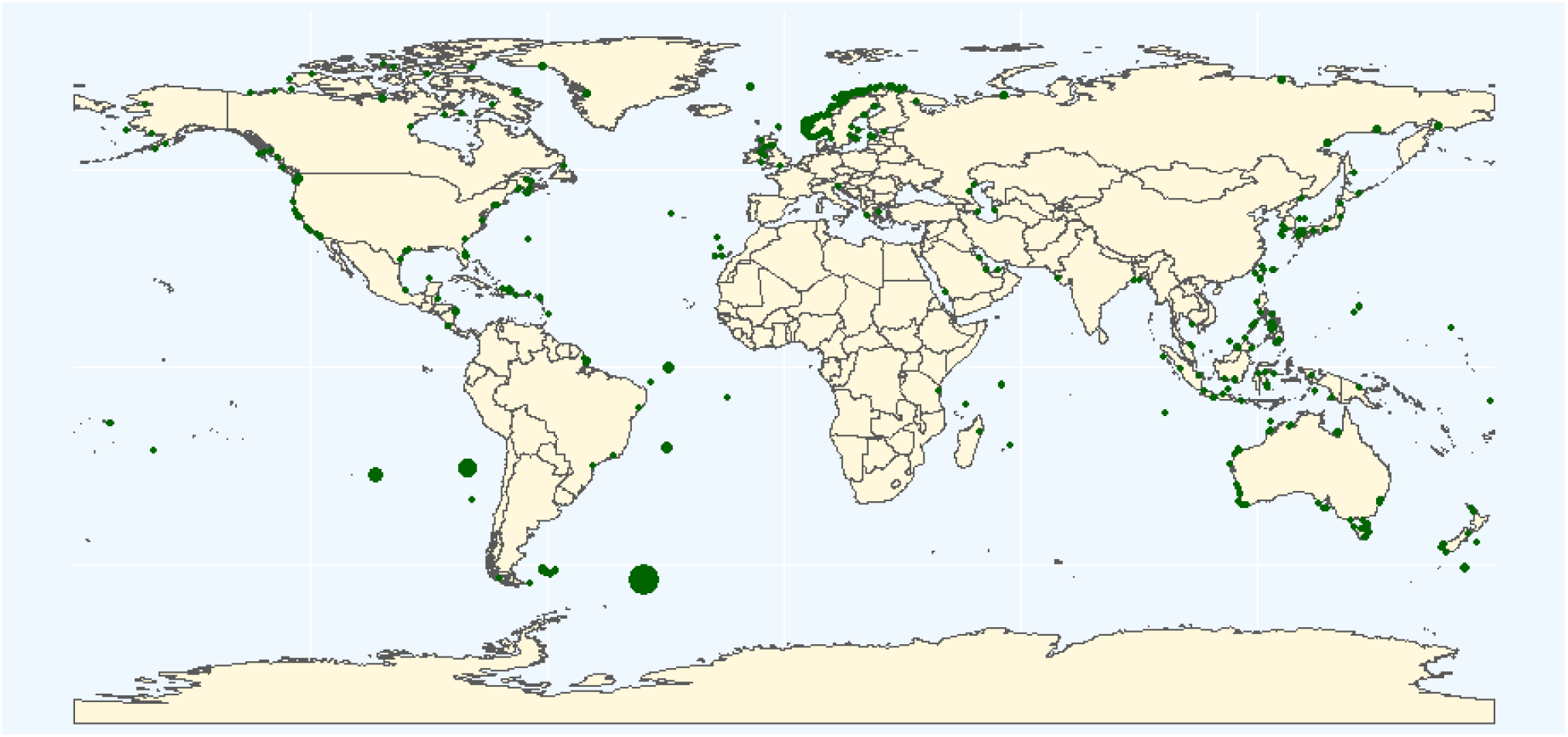
Isolated marine reserves that are not connected to any others with a PLD of 40 days. Dots show the relative surface area of each reserve.

### 3.2 Relationship between country protection coverage and connectedness

While networks were designated without directionality, the probability matrix does show which reserves act as sources and sinks. Combining reserves by country provides an overview of the hydrodynamics that connect countries, as opposed to individual reserves. This explains the flow of resources and shows which countries most benefit from incoming larval dispersal, and which countries can primarily serve as a source of larvae. Countries that act only as sources or sinks tend to control few marine reserves and, therefore, have few overall connections. For example, the Seychelles (8 reserves in total) have 3 outgoing connections but no incoming connections, while the Dominican Republic (2 reserves total) has 5 incoming connections but only 1 outgoing. Countries with more reserves show better connectivity and have a more equal ratio of incoming to outgoing connections. The USA is a good model for high connectivity with a good incoming to outgoing ratio: it manages both a large area of reserves and a high number of reserves, in total it has 186 reserves with 19 external connections, 11 of which are incoming and 8 are outgoing. Norway is also an extreme case, with many more reserves than any other country, but low surface coverage. Norway is also not highly connected to other countries, with external connections to only three countries despite controlling 610 reserves.

Overall, countries with more and larger marine reserves tend to be more connected to other countries (Figure 3). Both the number and size of reserves relate to the external connectivity a country has, but the size of reserves has a slightly stronger but still not significant relationship to the external connectivity than number of reserves has. Some countries have an unexpectedly high number of connections given their number and size of reserves, such as the US Virgin Islands, which has few reserves but many connections and overall a relatively even distribution of incoming to outgoing connections.

**Figure 3:**
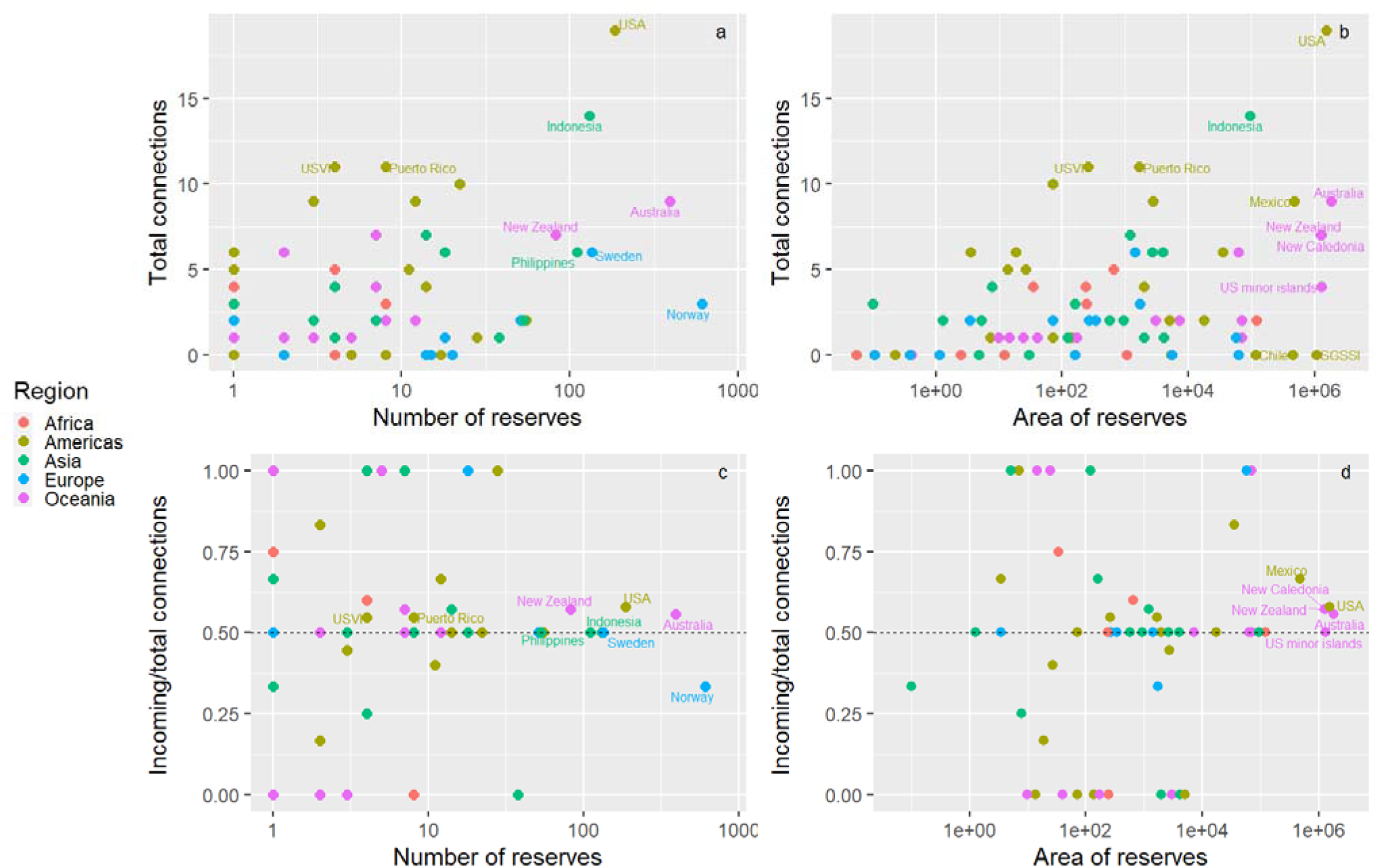
How the MRs of each country relate to that country’s hydrological connections to other countries, including how the number (a) and area (b) of MRs of one country relate to the total connections to all other countries within the same ecological network. The percent of connections in each country that are incoming only is show vs the number of reserves in (c) and vs the area of reserves in (d). Points are countries and point colors are geographic regions.

The number of connections to other countries’ marine reserves is more dependent on the total area of reserves each country controls, rather than the number of reserves (Figure 3). However, countries with a high number of reserves, as well as a high number of external connections are more likely to have a high human development index, have a low fisheries dependency score, and have a high environmental performance index (Figure 4).

**Figure 4:**
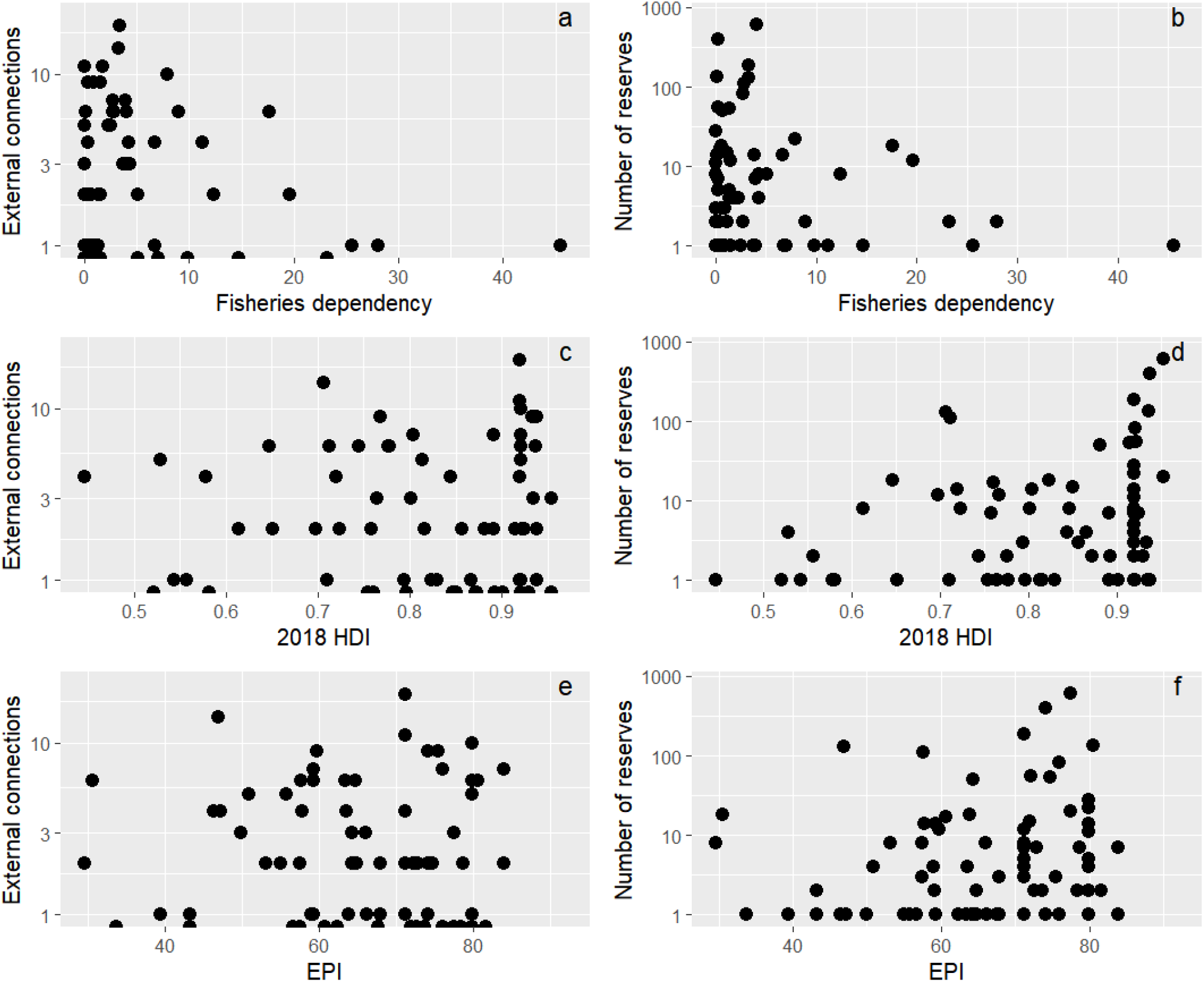
the number of external connections and the number of reserves controlled relate to the social, economic, and cultural characteristics of the countries that manage the reserves. External connections as a function of a) fisheries dependency, c) human development index and e) environmental performance index. Total number of reserves controlled by a country as a function of b) fisheries dependency, d) human development index, and f) environmental performance index.

## 4 Discussion

Overall, we found that the majority of marine reserves (73.4%) are connected to at least one other reserve, and of these networks more are very well connected in a large (>20 MR) network (60.7%). Of the seven largest networks that span multiple countries, few have diverse social and political indicators, and tend to be managed by countries with similar economic indicators. Our main results show that: 1) while most marine reserves across the globe are connected to other reserves, many are connected in large networks of more than 20 separate reserves that span multiple diverse countries, 2) isolated reserves make up a significant portion of total reserves (26.6%) and are present across the world, 3) and connections between marine reserves are directional, implying that some of them, and countries that control them, are likely to act more as sources or as sinks.

Here, we find that the majority of the highly connected networks are managed by multiple countries, meaning successful management of most of the world’s largest ecological networks will require coordination between countries. Countries with a high human development index have the most connections because they have the most reserves, as previously recognized (Marinesque et al. 2012). Countries with similar sociopolitical characteristics have fewer barriers to coordination; for example, Network 2 is split between Norway and Sweden, which both have a high human development index, a strong focus on the environment, and low fisheries dependency, as well as sharing a border. Canada and the United States also share a large network, as do Australia and New Zealand. These networks are likely to have an immediate advantage over networks with divergent cultural or socioeconomic attributes in terms of management and communication (Cámara-Leret 2019, Wang 2015, Putnam 2008). Over time however, higher levels of diversity have been shown to create better societies, as long as there is a common goal (Page 2008), so while diverse decision maker groups may prove to have lower levels of social trust in the first stages of cooperation, over the longer-term diverse networks could prove to be more ecologically efficient (Wang 2015). In addition, networks with a diverse range of EPIs might bolster the protection efforts of the included countries with lower EPIs with an overall positive effect.

Though connected networks are more prevalent, isolated reserves are well represented around the world, both in places with large networks and without. Isolated reserves near large networks are likely to be smaller, but make for good candidates to connect to the nearby large networks in the future. Many isolated large reserves in the open ocean are in highly diverse areas such as the Southern Atlantic. Isolated reserves are in danger of slower recovery after catastrophic events, and in having lower gene diversity. Identifying sites that are highly connected to, but not currently part of, a marine reserve or protected area is another good way to find new sites to extent existing networks of protected areas to achieve the ambitious goals of the post-2020 biodiversity framework (Magris et al 2018).

Many countries have asymmetric connectivity, and act primarily as either sources or sinks. As the number of reserves managed by a country increases, so does the symmetry. This is probably a sampling effect, caused by the fact that increasing the number of reserves will also increase the chances of including both source and sink reserves. Not only is the level of connectivity important, but the direction of the connectivity is also. Effectively connected networks can only be created if the importance of the direction of connection is taken into consideration by the managing body of the network.

Larval dispersal has the potential to reach up to hundreds of kilometers outside of reserves based on biophysical models, and recruitment from external sources can be a valuable component of recovery and persistence of coral reefs, with some reefs acting as sources for larvae and other more connected reserves acting as stepping-stones aiding in dispersal (Roberts 2017, Alvarez-Romero 2018). Larval dispersal is however mediated by larval behavior such as active swimming and orientation, and can result in different patterns of marine reserve connectivity compared to those identified using hydrodynamic connectivity (Faillettaz et al 2018). The identity and size of the marine reserve networks identified here might therefore be different when considered under the lens of marine larval dispersal.

These results demonstrate the need for more research into how different countries that share an ecological network interact regarding management measures. While this work gives a snapshot of similarities between countries, it does not delve into the actual interactions between the countries with regards to marine management and how external connectivity is managed. Identifying key areas of connectivity and ensuring convergent management styles to protect these areas would help countries better protect their marine resources, help vulnerable populations that depend on the sea for their livelihoods, and help mitigate some of the impacts of anthropogenic climate change.

## 6 Author Contributions

JM, DM, and MA conceived of the study, LV and NB contributed to analysis, JM wrote the first draft, and all authors commented on the manuscript.

## 9 Data Availability Statement

The datasets analyzed for this study can be found in the Protected Planet database [https://www.protectedplanet.net/en/thematic-areas/marine-protected-areas].

## References

1. Álvarez-Romero, J.G., Munguía-Vega, A., Beger, M., del Mar Mancha-Cisneros, M., Suárez-Castillo, A.N., Gurney, G.G., Pressey, R.L., Gerber, L.R., Morzaria-Luna, H.N., Reyes-Bonilla, H., Adams, V.M., Kolb, M., Graham, E.M., VanDerWal, J., Castillo-López, A., Hinojosa-Arango, G., Petatán-Ramírez, D., Moreno-Baez, M., Godínez-Reyes, C.R. & Torre, J. (2018). Designing connected marine reserves in the face of global warming. Glob. Chang. Biol., 24, e671–e691.

2. Andrello, M., Guilhaumon, F., Albouy, C., Parravicini, V., Scholtens, J., Verley, P., Barange, M., Sumaila, U.R., Manel, S. & Mouillot, D. (2017). Global mismatch between fishing dependency and larval supply from marine reserves. Nat. Commun., 8.

3. Ban, N.C., Gurney, G.G., Marshall, N.A., Whitney, C.K., Mills, M., Gelcich, S., Bennett, N. J., Meehan, M.C., Butler, C., Ban, S., Tran, T.C., Cox, M.E. & Breslow, S.J. (2019). Well-being outcomes of marine protected areas. Nat. Sustain., 2, 524–532.

4. Bivand R, Rundel C (2017) rgeos: Interface to geometry engine-open source (‘geos’). R package version 0.3-26. https://CRAN.R-project.org/package=rgeos

5. Bodin, Ö., Robins, G., Mcallister, R.R.J.J., Guerrero, A.M., Tengö, M., Lubell, M., Crona, B., Tengö, M. & Lubell, M. (2016). Theorizing benefits and constraints in collaborative environmental governance: A transdisciplinary social-ecological network approach for empirical investigations. Ecol. Soc., 21.

6. Booth, D.J. & Parkinson, K. (2011). Pelagic larval duration is similar across 23° of latitude for two species of butterflyfish (Chaetodontidae) in eastern Australia. Coral Reefs, 30, 1071–1075.

7. Cámara-Leret, R., Fortuna, M.A. & Bascompte, J. (2019). Indigenous knowledge networks in the face of global change. Proc. Natl. Acad. Sci. U. S. A., 116, 9913–9918.

8. Cinner, J.E., Maire, E., Huchery, C., Aaron MacNeil, M., Graham, N.A.J., Mora, C., McClanahan, T.R., Barnes, M.L., Kittinger, J.N., Hicks, C.C., D’Agata, S., Hoey, A.S., Gurney, G.G., Feary, D.A., Williams, I.D., Kulbicki, M., Vigliola, L., Wantiez, L., Edgar, G.J., Stuart-Smith, R.D., Sandin, S.A., Green, A., Hardt, M.J., Beger, M., Friedlander, A.M., Wilson, S.K., Brokovich, E., Brooks, A.J., Cruz-Motta, J.J., Booth, D.J., Chabanet, P., Gough, C., Tupper, M., Ferse, S.C.A., Rashid Sumaila, U., Pardede, S. & Mouillot, D. (2018). Gravity of human impacts mediates coral reef conservation gains. Proc. Natl. Acad. Sci. U. S. A., 115, E6116–E6125.

9. Costello, M.J. & Ballantine, B. (2015). Biodiversity conservation should focus on no-take Marine Reserves: 94% of Marine Protected Areas allow fishing. Trends Ecol. Evol., 30, 507–509.

10. Csardi G, Nepusz T (2006). “The igraph software package for complex network research.” InterJournal, Complex Systems, 1695. https://igraph.org.

11. CTI-CFF Regional Secretariat (2016). Annual Activities Report 2015-2016. coraltriangleinitiative.org

12. D. Putnam’s “E Pluribus Unum: Diversity and community in the twenty-first century the 2006 Johan Skytte Prize Lecture.” Hous. Policy Debate, 19, 207.

13. Dinerstein, E., Vynne, C., Sala, E., Joshi, A.R., Fernando, S., Lovejoy, T.E., Mayorga, J., Olson, D., Asner, G.P., Baillie, J.E.M., Burgess, N.D., Burkart, K., Noss, R.F., Zhang, Y.P., Baccini, A., Birch, T., Hahn, N., Joppa, L.N. & Wikramanayake, E. (2019). A Global Deal for Nature: Guiding principles, milestones, and targets. Sci. Adv., 5, 1–18.

14. Duarte, C.M., Agusti, S., Barbier, E., Britten, G.L., Castilla, J.C., Gattuso, J.P., Fulweiler, R.W., Hughes, T.P., Knowlton, N., Lovelock, C.E., Lotze, H.K., Predragovic, M., Poloczanska, E., Roberts, C. & Worm, B. (2020). Rebuilding marine life. Nature, 580, 39–51.

15. Eberhard, David M., Gary F. Simons, and Charles D. Fennig (eds.). (2019). Ethnologue: Languages of the World. Twenty-second edition. Dallas, Texas: SIL International.

16. Edgar, G.J., Stuart-Smith, R.D., Willis, T.J., Kininmonth, S., Baker, S.C., Banks, S., Barrett, N.S., Becerro, M.A., Bernard, A.T.F., Berkhout, J., Buxton, C.D., Campbell, S.J., Cooper, A.T., Davey, M., Edgar, S.C., Försterra, G., Galván, D.E., Irigoyen, A.J., Kushner, D.J., Moura, R., Parnell, P.E., Shears, N.T., Soler, G., Strain, E.M.A. & Thomson, R.J. (2014). Global conservation outcomes depend on marine protected areas with five key features. Nature, 506, 216–220.

17. Faillettaz, R., Paris, C. B., and Irisson, J. O. (2018). Larval fish swimming behavior alters dispersal patterns from marine protected areas in the North-Western Mediterranean Sea. Front. Mar. Sci. 5, 1–12. doi:10.3389/fmars.2018.00097.

18. Goetze, J.S., Wilson, S., Radford, B., Fisher, R., Langlois, T.J., Monk, J., Knott, N.A., Malcolm, H., Currey-, L.M., Daniel, R., David, I., Barrett, N., Babcock, R.C., Bosch, N.E., Brock, D., Claudet, J., Clough, J., Fairclough, D. V, Heupel, M.R., Holmes, T.H., Huveneers, C., Jordan, A.R., Mclean, D., Meekan, M., Miller, D., Newman, S.J., Rees, M.J., Roberts, K.E., Saunders, B.J., Speed, C.W., Eric, J.T., Sasha, T., Corey, K.W. & Harvey, E.S. (2021). Increased connectivity and depth improve the effectiveness of marine reserves, 3432–3447.

19. Graham, E.M., Baird, A.H. & Connolly, S.R. (2008). Survival dynamics of scleractinian coral larvae and implications for dispersal. Coral Reefs, 27, 529–539.

20. Hughes, T.P., Kerry, J.T., Connolly, S.R., Baird, A.H., Eakin, C.M., Heron, S.F., Hoey, A.S., Hoogenboom, M.O., Jacobson, M., Liu, G., Pratchett, M.S., Skirving, W. & Torda, G. (2019). Ecological memory modifies the cumulative impact of recurrent climate extremes. Nat. Clim. Chang., 9, 40–43.

21. Jaspers, C., Huwer, B., Antajan, E., Hosia, A., Hinrichsen, H.H., Biastoch, A., Angel, D., Asmus, R., Augustin, C., Bagheri, S., Beggs, S.E., Balsby, T.J.S., Boersma, M., Bonnet, D., Christensen, J.T., Dänhardt, A., Delpy, F., Falkenhaug, T., Finenko, G., Fleming, N.E.C., Fuentes, V., Galil, B., Gittenberger, A., Griffin, D.C., Haslob, H., Javidpour, J., Kamburska, L., Kube, S., Langenberg, V.T., Lehtiniemi, M., Lombard, F., Malzahn, A., Marambio, M., Mihneva, V., Møller, L.F., Niermann, U., Okyar, M.I., Özdemir, Z.B., Pitois, S., Reusch, T.B.H., Robbens, J., Stefanova, K., Thibault, D., van der Veer, H.W., Vansteenbrugge, L., van Walraven, L. & Woźniczka, A. (2018). Ocean current connectivity propelling the secondary spread of a marine invasive comb jelly across western Eurasia. Glob. Ecol. Biogeogr., 27, 814–827.

22. Jones, G.P., Almany, G.R., Russ, G.R., Sale, P.F., Steneck, R.S., Van Oppen, M.J.H. & Willis, B.L. (2009). Larval retention and connectivity among populations of corals and reef fishes: History, advances and challenges. Coral Reefs, 28, 307–325.

23. Jönsson, B.F. & Watson, J.R. (2016). The timescales of global surface-ocean connectivity. Nat. Commun., 7, 1–6.

24. Lett, C., Verley, P., Mullon, C., Parada, C., Brochier, T., Penven, P. & Blanke, B. (2008). A Lagrangian tool for modelling ichthyoplankton dynamics. Environ. Model. Softw., 23, 1210–1214.

25. Magris, R.A., Andrello, M., Pressey, R.L., Mouillot, D., Dalongeville, A., Jacobi, M.N. & Manel, S. (2018). Biologically representative and well-connected marine reserves enhance biodiversity persistence in conservation planning. Conserv. Lett., 11, 1–10.

26. Manel, S., Loiseau, N., Andrello, M., Fietz, K., Goñi, R., Forcada, A., Lenfant, P., Kininmonth, S., Marcos, C., Marques, V., Mallol, S., Pérez-Ruzafa, A., Breusing, C., Puebla, O. & Mouillot, D. (2019). Long-Distance Benefits of Marine Reserves: Myth or Reality? Trends Ecol. Evol., 34, 342–354.

27. Marinesque, S., Kaplan, D. M., and Rodwell, L. D. (2012). Global implementation of marine protected areas: Is the developing world being left behind? Mar. Policy 36, 727–737. doi:10.1016/j.marpol.2011.10.010.

28. O’Hara, C. C., Frazier, M., and Halpern, B. S. (2021). At-risk marine biodiversity faces extensive, expanding, and intensifying human impacts. Science (80-.). 372, 84–87. doi:10.1126/science.abe6731.

29. O’Leary, B.C., Ban, N.C., Fernandez, M., Friedlander, A.M., García-Borboroglu, P., Golbuu, Y., Guidetti, P., Harris, J.M., Hawkins, J.P., Langlois, T., McCauley, D.J., Pikitch, E.K., Richmond, R.H. & Roberts, C.M. (2018). Addressing Criticisms of Large-Scale Marine Protected Areas. Bioscience, 68, 359–370.

30. Ohayon, S., Granot, I. & Belmaker, J. (n.d.). marine protected areas.

31. Page SE. (2008). How the Power of Diversity Creates Better Groups, Firms, Schools, and Societies. Princeton University Press.

32. Roberts, C.M., O’Leary, B.C., Mccauley, D.J., Cury, P.M., Duarte, C.M., Lubchenco, J., Pauly, D., Sáenz-Arroyo, A., Sumaila, U.R., Wilson, R.W., Worm, B. & Castilla, J.C. (2017). Marine reserves can mitigate and promote adaptation to climate change. Proc. Natl. Acad. Sci. U. S. A., 114, 6167–6175.

33. Sala, E. & Giakoumi, S. (2018). No-take marine reserves are the most effective protected areas in the ocean. ICES J. Mar. Sci., 75, 1166–1168.

34. Smith, T.W., Kim, J. & Son, J. (2017). Public Attitudes toward Climate Change and Other Environmental Issues across Countries. Int. J. Sociol., 47, 62–80.

35. Treml, E.A., Fidelman, P.I.J., Kininmonth, S., Ekstrom, J.A. & Bodin, Ö. (2015). Analyzing the (Mis) Fit Between Institutional and Ecological Networks. Glob. Environ. Chang., 31, 263–271.

36. Turnbull, J.W., Johnston, E.L. & Clark, G.F. (2021). Evaluating the social and ecological effectiveness of partially protected marine areas, 35, 921–932.

37. UNDP. 2018. 2018 Statistical Update: Human Development Indices and Indicators. New York. http://hdr.undp.org/en/content/human-development-indices-indicators-2018 (accessed January 2019)

38. UNEP-WCMC and IUCN. Protected planet: The World Database on Protected Areas (WDPA). UNEP-WCMC and IUCN, Cambridge, UK. Available from www.protectedplanet.net (accessed January 2019).

39. Wang, C. & Steiner, B. (2015). Can Ethno-Linguistic Diversity Explain Cross-Country Differences in Social Capital?: A Global Perspective. Econ. Rec., 91, 338–366.

40. Wendling, Z. A., Emerson, J. W., de Sherbinin, A., Esty, D. C., et al. (2020). 2020 Environmental Performance Index. New Haven, CT: Yale Center for Environmental Law & Policy. epi.yale.edu

