## Supplemental information for "Global connectivity and networks of marine reserves"

Supplementary Material

### 1 Supplementary Data

Methods

From the initial Protected Planet dataset containing 9,600 MPAs, we retained only MPAs that met the following filtering criteria. First, we removed MPAs located entirely on land using the land ecoregion of the world polygon (www.worldwildlife.org/biomes). Second, we retained only the MRs that are either in IUCN category Ia or Ib or designated as at least partially no-take, since only marine reserves provide widely recognized ecological benefits (Edgar et al. 2014, 2018, Strain et al. 2018, Cinner et al. 2018).

Sea surface current velocities were obtained by the Copernicus Marine Environment Monitoring Service ocean physical reanalysis GLOBAL_REANALYSIS_PHY_001_030-TDS. The horizontal resolution of the model is 1/12° and the temporal resolution of stored data is one day at a depth of one meter, and runs from 180°W–180°E, 80°S–80°N. We used data from Jan 01 2008 to March 03 2017, the most recent output at the time of the study. Hydrodynamic dispersal simulations were performed with Ichthyop 3.3 (Lett et al. 2008). Ten thousand virtual particles were released in the centroid of each MR at the midpoint of each season (2nd February, 5th May, 6th August and 11th November) in each of the 10 years between 2008–2017, for a total of 883,600,000 released particles. Simulation had a time step of 3,600 s (1 h), which is sufficiently short for a particle to not cross more than one boundary of hydrodynamic cells in one step. Particle positions were recorded every 5 days, up to one year (365 days). Particles were subject to passive dispersal only. Advection was simulated using a Runge-Kutta 4^th^ order numerical scheme. Horizontal diffusion was applied via a random walk to account for sub-grid-scale hydrodynamics following Peliz et al. (2007), with a horizontal diffusion coefficient *K* = *ε*^1/3^*l*^4/3^, where *ε* = 10^-9^ m^2^s^-3^ is the constant turbulent dissipation rate (Monin & Ozmidov, 1982) and *l* is length of the grid cell.

### 2 Supplementary Figures and Tables

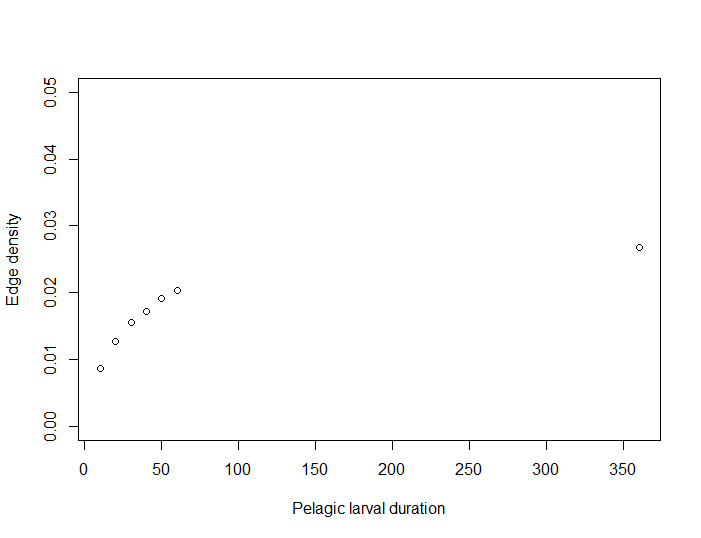

Supplementary Figure 1. Effect of pelagic larval duration on the edge density (ratio of number of realized edges to the number of potential edges). While the pelagic larval duration (PLD) does influence the size of the networks with longer PLDs leading to larger networks with more reserves, this relationship appears to stabilize around 40 days. In addition, the edge density of networks (which is the ratio of number of realized edges to the number of potential edges) also saturates at ~0.27 indicating that most connections are created by 60 days.

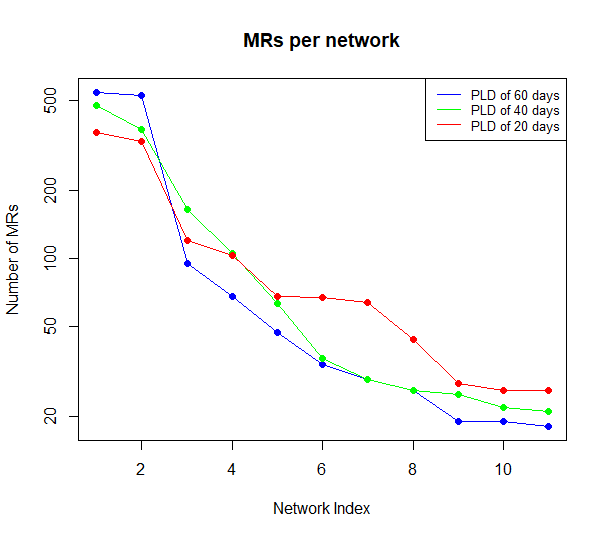

Supplementary Figure 2. Size of the 11 largest networks for the three different pelagic larval durations. The number of marine reserves in each network, when ranked from largest to smallest network, also followed similar patterns between 20, 40, and 60 days (Appendix 3). For each PLD analyzed, around 26% of the reserves are not included in any network.

Supplementary Table 1. The composition of the largest networks of marine reserves, ordered by number of reserves

| ^Reserve Ranking^ | ^Number of Reserves^ | ^Rank of Area^ | ^Area (kmsq)^ | ^Number of Languages^ | ^Number of Countries^ |
| --- | --- | --- | --- | --- | --- |
| ^1^ | ^476^ | ^1^ | ^3859863^ | ^3^ | ^5^ |
| ^2^ | ^374^ | ^43^ | ^1653^ | ^2^ | ^2^ |
| ^3^ | ^165^ | ^41^ | ^2293^ | ^3^ | ^3^ |
| ^4^ | ^105^ | ^19^ | ^15068^ | ^2^ | ^2^ |
| ^5^ | ^63^ | ^151^ | ^14^ | ^1^ | ^1^ |
| ^6^ | ^36^ | ^69^ | ^618^ | ^3^ | ^5^ |
| ^7^ | ^29^ | ^28^ | ^5754^ | ^1^ | ^1^ |
| ^8^ | ^26^ | ^155^ | ^13^ | ^1^ | ^1^ |
| ^9^ | ^25^ | ^18^ | ^21417^ | ^2^ | ^4^ |
| ^10^ | ^22^ | ^13^ | ^38728^ | ^1^ | ^1^ |
| ^11^ | ^21^ | ^4^ | ^356779^ | ^1^ | ^2^ |

Supplementary references

1. Cinner, J.E., Maire, E., Huchery, C., Aaron MacNeil, M., Graham, N.A.J., Mora, C., McClanahan, T.R., Barnes, M.L., Kittinger, J.N., Hicks, C.C., D’Agata, S., Hoey, A.S., Gurney, G.G., Feary, D.A., Williams, I.D., Kulbicki, M., Vigliola, L., Wantiez, L., Edgar, G.J., Stuart-Smith, R.D., Sandin, S.A., Green, A., Hardt, M.J., Beger, M., Friedlander, A.M., Wilson, S.K., Brokovich, E., Brooks, A.J., Cruz-Motta, J.J., Booth, D.J., Chabanet, P., Gough, C., Tupper, M., Ferse, S.C.A., Rashid Sumaila, U., Pardede, S. & Mouillot, D. (2018). Gravity of human impacts mediates coral reef conservation gains. *Proc. Natl. Acad. Sci. U. S. A.*, 115, E6116–E6125.
2. Edgar, G.J., Stuart-Smith, R.D., Willis, T.J., Kininmonth, S., Baker, S.C., Banks, S., Barrett, N.S., Becerro, M.A., Bernard, A.T.F., Berkhout, J., Buxton, C.D., Campbell, S.J., Cooper, A.T., Davey, M., Edgar, S.C., Försterra, G., Galván, D.E., Irigoyen, A.J., Kushner, D.J., Moura, R., Parnell, P.E., Shears, N.T., Soler, G., Strain, E.M.A. & Thomson, R.J. (2014). Global conservation outcomes depend on marine protected areas with five key features. *Nature*, 506, 216–220.
3. Edgar, G.J., Ward, T.J. & Stuart-Smith, R.D. (2018). Rapid declines across Australian fishery stocks indicate global sustainability targets will not be achieved without an expanded network of ‘no-fishing’ reserves. *Aquat. Conserv. Mar. Freshw. Ecosyst.*, 28, 1337–1350.
4. Lett, C., Verley, P., Mullon, C., Parada, C., Brochier, T., Penven, P. & Blanke, B. (2008). A Lagrangian tool for modelling ichthyoplankton dynamics. *Environ. Model. Softw.*, 23, 1210–1214.
5. Monin, A.S., Ozmidov, R.V., 1981. Ocean Turbulence. Gidrometeoizdat, Leningrad.
6. Peliz, A., Marchesiello, P., Dubert, J. & Marta-almeida, M. (2007). A study of crab larvae dispersal on the Western Iberian Shelf : Physical processes, 68, 215–236.
7. Strain, E.M.A., Edgar, G.J., Ceccarelli, D., Stuart-Smith, R.D., Hosack, G.R. & Thomson, R.J. (2019). A global assessment of the direct and indirect benefits of marine protected areas for coral reef conservation. *Divers. Distrib.*, 25, 9–20.
